# Erythroid precursors and progenitors suppress adaptive immunity and get invaded by SARS-CoV-2

**DOI:** 10.1101/2020.08.18.255927

**Authors:** Shima Shahbaz, Lai Xu, Mohammad Osman, Wendy Sligl, Justin Shields, Michael Joyce, Lorne Tyrrell, Olaide Oyegbami, Shokrollah Elahi

**Affiliations:** School of Dentistry, Division of Foundational Sciences, University of Alberta, Edmonton, T6G2E1, AB, Canada; Department of Medicine, University of Alberta, Edmonton, T6G2E1, AB, Canada; Department of Critical Care Medicine, University of Alberta, Edmonton, T6G2E1, AB, Canada; Division of Infectious Diseases, University of Alberta, Edmonton, T6G2E1, AB, Canada; Department of Medical Microbiology and Immunology, University of Alberta, Edmonton, T6G2E1, AB, Canada; Li Ka Shing Institute of Virology, University of Alberta, Edmonton, T6G2E1, AB, Canada; Department of Medical Oncology, Faculty of Medicine and Dentistry, University of Alberta, Edmonton, T6G2E1, AB, Canada

**Keywords:** COVID-19, CD71+ erythroid cells, RBCs, SARS-CoV-2, dexamethasone

## Abstract

SARS-CoV-2 infection is associated with lower blood oxygen levels even in patients without hypoxia requiring hospitalization. This discordance illustrates the need for a more unifying explanation as to whether SARS-CoV-2 directly or indirectly affects erythropoiesis. Here we show significantly enriched CD71+ erythroid precursors/progenitors in the blood circulation of COVID-19 patients that have distinctive immunosuppressive properties. A subpopulation of abundant erythroid cells, CD45+CD71+cells, co-express ACE2, TMPRSS2, CD147, CD26 and these can be infected with SARS-CoV-2. In turn, pre-treatment of erythroid cells with dexamethasone significantly diminished ACE2/TMPRSS2 expression and subsequently reduced their infectivity with SARS-CoV-2. Taken together, pathological abundance of erythroid cells might reflect stress erythropoiesis due to the invasion of erythroid progenitors by SARS-CoV-2. This may provide a novel insight into the impact of SARS-CoV-2 on erythropoiesis and hypoxia seen in COVID-19 patients.

## Introduction

The COrona VIrus Disease 2019 (COVID-19) caused by severe acute respiratory syndrome coronavirus 2 (SARS-CoV-2) has resulted in global crisis. SARS-CoV-2 infection manifests as a spectrum from asymptomatic or mild symptoms to moderate and severe disease^1^. A subgroup will become critically ill and develop acute respiratory distress syndrome (ARDS), a clinical phenomenon characterized by the development of bilateral infiltrates and hypoxemia^2^, often accompanied with septic shock and organ falure^3,4^.

The pathogenesis of SARS-CoV2 is being delineated rapidly, however the causes of hypoxia, have remained elusive. It has been shown that SARS-CoV-2 infection is initiated by the viral surface spike glycoprotein (S protein)^5^ binding to the angiotensin-converting enzyme-2 (ACE2) for cell entry^6^. Subsequently, the S protein gets cleaved by the transmembrane protease serin2 (TMPRSS2)^6^. It appears that SARS-CoV-2 not only gains initial entry through ACE2 but also downregulates cell surface ACE2 expression such that this enzyme cannot exert its protective role ^7^. Downregulation of ACE2 in the respiratory tract is linked to neutrophils infiltration in response to LPS^8^ and may result in angiotensin II accumulation and lung injury as has been reported in animal models of respiratory virus infections^9,10^. In addition to the respiratory tract, ACE2 expression has been reported in intestinal epithelial cells, endothelial cells, renal tubules, cerebral neurons and possibly immune cells (e.g. alveolar monocytes/macrophages) ^11^. Reduced numbers of T, B and NK cells in the peripheral blood of COVID-19 patients has been reported, especially in those with severe disease^4,12,13^. In spite of elevated levels of granulocyte macrophage colony stimulating factor a decline in the proportion of monocytes, eosinophils and basophils has been reported^12^. In contrast to what occurs in peripheral blood, higher neutrophil recruitment in the lungs has been associated with disease severity^12^. Despite the frequency of hypoxia, the impact of SARS-CoV-2 infection on erythropoiesis has received limited attention. Preliminary modeling reports have suggested that SARS-CoV-2 may inhibit heme metabolism and induce hemoglobin denaturation^14^. As such, hemoglobin alteration may compromise oxygen-carrying capacity of red blood cells (RBCs) in COVID-19 patients resulting in hypoxia. The entry receptor ACE2, has been confirmed in RBCs by proteomic studies^15^. This suggests that RBCs might be targeted by SARS-CoV-2 virus, although they cannot support viral replication. RBCs can be directly invaded by pathogens (e.g. in malaria), resulting in hemolysis ^16^. In support of this concept, structural protein damage and changes in RBC membrane lipids have been reported in COVID-19 patients^17^. In addition to ACE2, SARS-CoV-2 invades host cells via CD147^18^, a known RBC receptor for *Plasmodium falciparum^19^*. Lastly, CD26 was reported to interact with SARS-CoV-2 spike^20^, which is involved in stress hematopoiesis^21^. In light of the above, it is possible that SARS-CoV-2 directly or indirectly invades RBCs. Hence, depletion of RBCs by SARS-CoV-2 may result in stress erythropoiesis as a compensatory mechanism to meet the oxygen supply, resulting in the abundance of erythroid precursors in the blood. Erythroid precursors are defined as CD71^+^ erythroid cells (CECs) co-expressing CD71 (the transferrin receptor) and CD235a (glycophorin A, erythroid lineage marker) in humans, and CD71 and TER119 in mice^22–24^. CECs are a heterogenous population of erythroid progenitors and precursors with a wide range of immunosuppressive and/or immunomodulatory properties ^25^. We and others have reported that CECs compromise innate and adaptive immune responses against infections and tumors due to their immunosuppressive properties ^4,26–30^. In addition, it has been shown that CECs can harbor infective HIV particles and the binding of HIV to CD235a mediates HIV trans-infection to CD4^+^ T cells^31^. However, whether these cells can be the target of SARS-CoV-2 remains to be explored.

In light of the above, we investigated the frequency and functionality of CECs in different groups of COVID-19 patients. We detected higher expression of ACE2, TMPRSS2 and CD147 on CECs compared to other immune cell lineages. We found that CECs can be infected with SARS-CoV-2 and infection can be partially inhibited by dexamethasone.

### Study population

Among 70 patients included in this study, 11 were critically ill patients (age 72.9±14.6 years) admitted to the Intensive Care Unit (ICU), whom we defined as having severe disease. Twenty three individuals (age 64.3±18.9) were hospitalized on a hospital ward > 5 days with moderate disease and the remaining 36 patients had mild disease requiring less than 5 days in hospital (age 61±17). ICU patients were older and 72.7% male (8/11) while non-ICU patients were 52.5% male (31/59). The mean age average for men and women were (67.7±14.5) and (62.8±22.6) respectively. Patient age ranged from17-95 years. Fifteen healthy individuals were recruited as negative controls (age 48±14.2).

### COVID-19 infection results in the expansion of CECs in the peripheral blood

The low oxygen saturation observed in COVID-19 patients^14^, suggested SARS-CoV-2 infection may have an effect on erythropoiesis. In this study, we observed that COVID-19 infection results in the expansion of CECs in the peripheral blood of patients compared to healthy controls (HCs). Of note, patients with severe COVID-19 disease had significantly higher percentages of CECs in peripheral blood compared to those with a moderate or mild disease (Fig. 1 A, 1B and Extended Data Fig.1A) and CECs were very low or absent in HCs (Fig. 1A and 1B). As the disease progressed over time, the CECs expanded in peripheral blood (Fig. 1C). We monitored the frequency of CECs over the entire disease course in three patients admitted to hospital. As shown in Fig. 1D and 1E, CECs expanded gradually after hospitalization but increased rapidly with the development of critical illness and eventually declined as patients no longer had detectable virus. In addition to CECs, we also observed an increase in the quantity of lighter weight RBCs (CD235a+CD71-cells) in the peripheral blood of patients while their presence in HCs was negligible (Extended Data Fig. 1B and C), again indicating the impact of COVID-19 infection on erythrocytes. We reasoned that the underlying mechanism for the expansion of CECs in COVID-19 patients might be related to dysregulated activity of hematopoietic stem and progenitor cells (HSPCs). We first examined IL-33 levels in plasma since elevated levels of IL-33 may inhibit the differentiation of CECs to RBCs ^32^. However, we did not observe any detectable level of IL-33 in our patients. Taken together, these observations suggest that SARS-CoV-2 infection influences erythropoiesis, resulting in the release of erythroid precursors and progenitors into the blood circulation.

**Fig 1.**
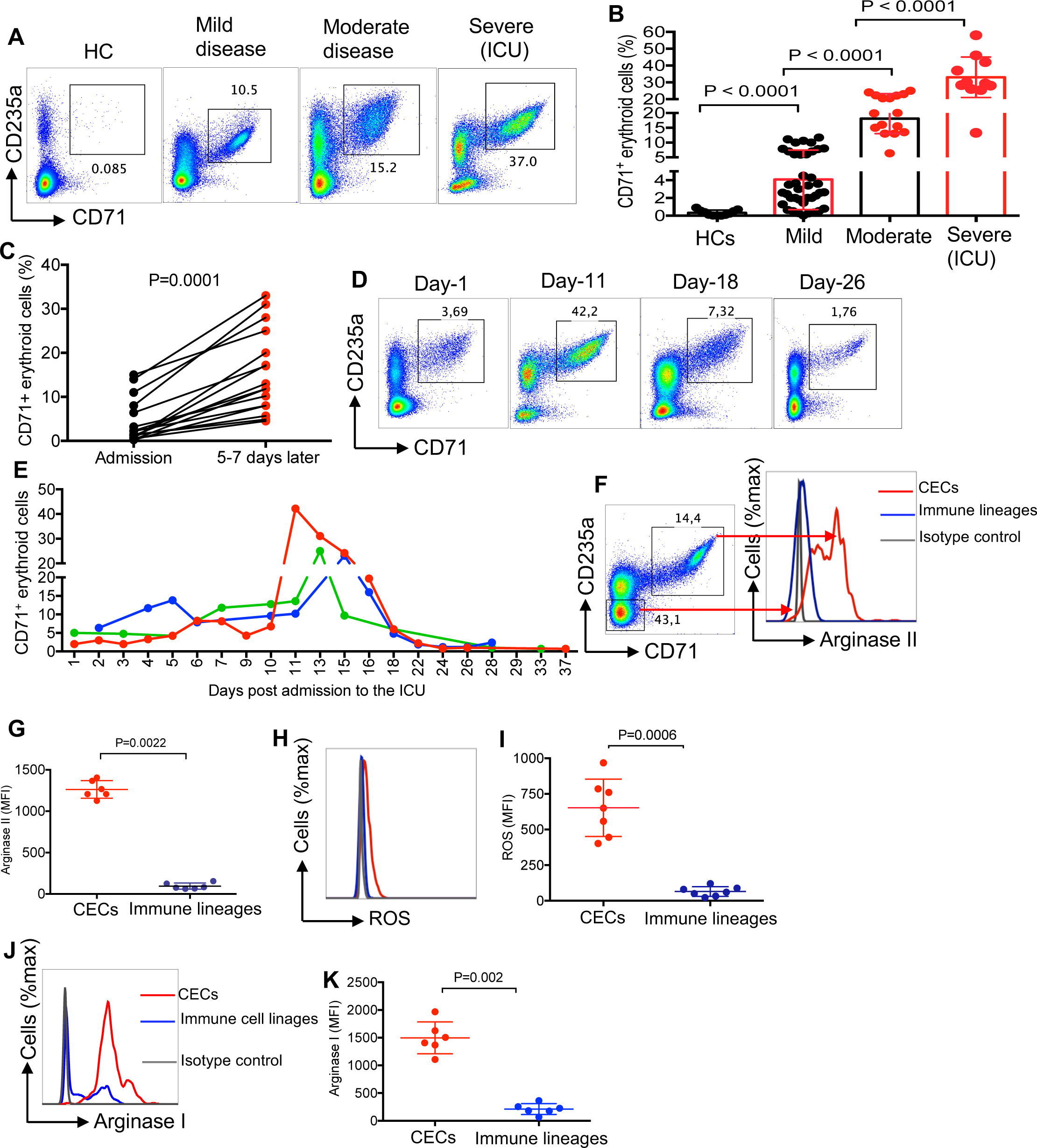
Expansion of CECs in COVID-19 patients is associated with the disease progression. (**A**) Representative flow cytometry plots, and (**B**) cumulative data of percentages of CECs in the PBMCs of COVID-19 patients versus healthy controls (HCs). (**C**) Data showing % CECs in PBMCs of patients at the time of admission to the hospital and 5-7 days later. (**D**) Representative plots, and (**E**) cumulative data of longitudinal changes in the frequency of CECs in 3 patients admitted to the ICU, each line represents a patient. (**F**) Representative plots, and (**G**) cumulative data of arginase-II expression in CECs compared to immune cell lineages. (**H**) Representative plots, and (**I**) cumulative data of ROS expression in CECs compared to other immune cell lineages in COVID-19 patients. (**J**) Representative plots, and (**K**) cumulative data of arginase-I expression in CECs compared to immune cell lineages in COVID-19 patients. Each point represents data from an individual/patient. Bar, mean ± one standard error.

### CECs express arginase II, arginase I, and reactive oxygen species (ROS) to mediate immunosuppression

Due to their immunomodulatory properties, the pathological abundance of CECs during disease progression can have immunological consequences ^22–23,25^. To better understand the biological properties of these expanded CECs in COVID-19 patients, we subjected them to immunological phenotyping. In contrast to other reports^30,33^, CECs in COVID-19 patients expressed negligible amount of PDL-1/PDL-2 but expressed the V-domain Immunoglobulin (Ig) Suppressor of T Cell Activation (VISTA) (Extended Data Fig. 1D). In agreement with our previous reports in other models ^24,26^, we found that CECs express significantly higher amounts of arginase II (Fig. 1F and 1G) and ROS (Fig. 1H and 1I) compared to other immune cell lineages, similar to what has been described for their counterparts in HIV^31^ and cancer^29^. For the very first time, we also detected expression of arginase I in CECs of COVID-19 patients (Fig. 1J and 1K). These observations guided us to investigate their immunosuppressive properties *in vitro*. CECs isolated from the peripheral blood mononuclear cells (PBMCs) that were > 95% pure (Extended Data Fig. 1E) were co-cultured with PBMCs at ratios of 1:1 or 1:2. The CECs significantly suppressed cytokine production (e.g. TNF-α and IFN-γ) by both CD4^+^ and CD8^+^ T cells when stimulated with anti-CD3/CD28 *in vitro* (Fig. 2A and 2B). CECs also impaired SARS-CoV-2 antigen-specific T cells when stimulated with overlapping peptide pools covering the main SARS-CoV-2 structural proteins-spike (S) and nucleocapsid (N) (Fig. 2C and 2D). Of note, antigen-specific response was dominated by TNF-αbut not IFN-γ(Fig. 2C and 2D). The CEC’s had a similar immunosuppressive effect on the capacity of CD8^+^ T cells to degranulate in response to viral peptide stimulation as measured using CD107a (Fig. 2E and Extended Data Fig. 1F). In agreement with previous reports in other models^29,34^, CECs significantly inhibited T cell proliferation following stimulation of PBMCs with SARS-CoV-2 peptides *in vitro* (Fig. 2F and 2G). This was supported *in vivo* by the negative correlation between the percentages of CECs, and CD3^+^ (Fig. 2H), CD4^+^ (Fig. 2I) and CD8^+^ T cells (Fig. 2J) in COVID-19 patients. We also observed an inverse correlation between CECs and the frequency of antibody secreting cells (plasmablasts) in COVID-19 (Extended Data Fig. 1G and 1H).

**Fig 2.**
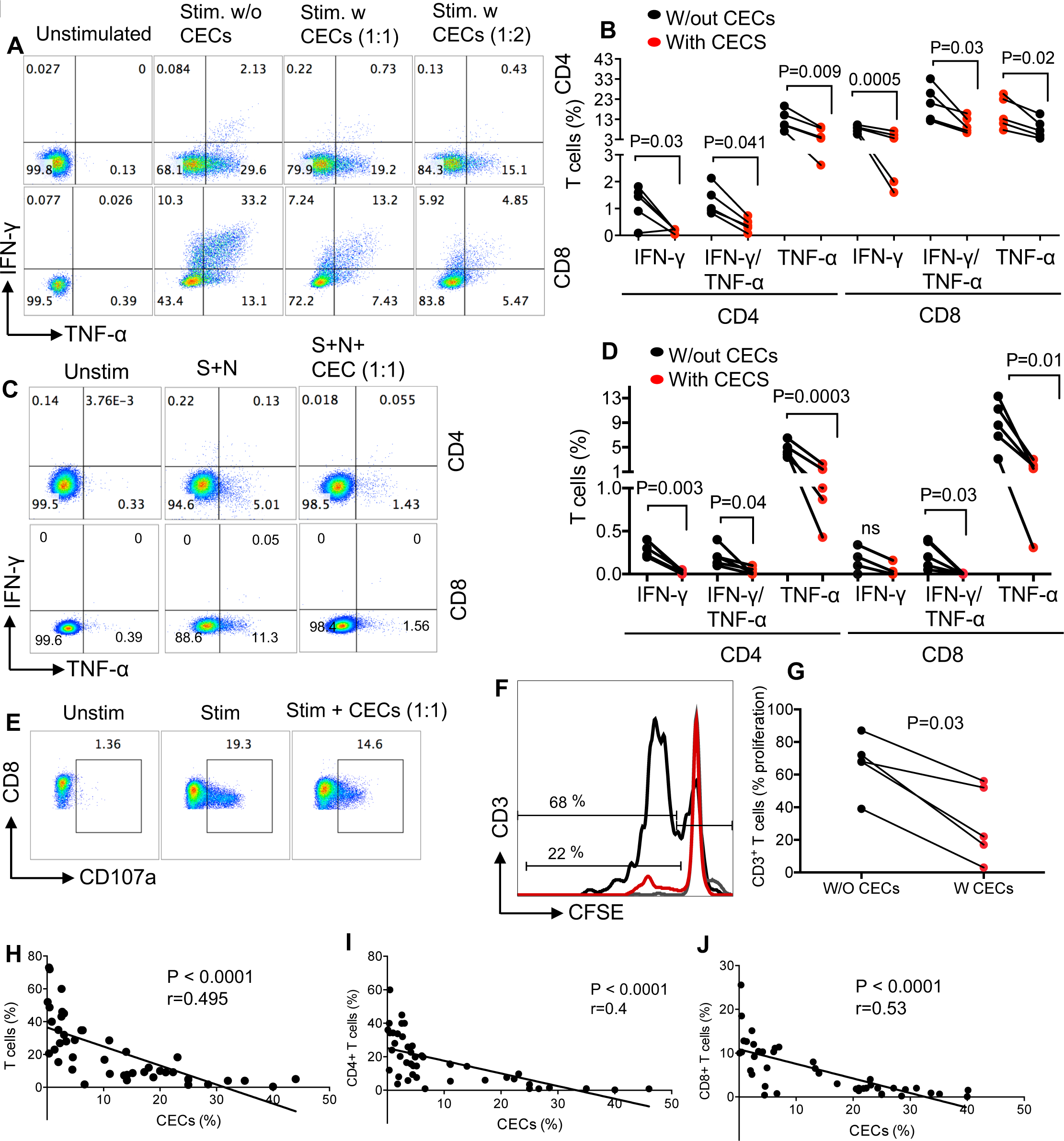
CECs exhibit immunosuppressive properties and their abundance may compromise adaptive immunity. (**A**) Representative plots, and (**B**) cumulative data of IFN-γand TNF-αexpression in CD4 and CD8 T following stimulation of PBMCs with anti-CD3/CD28 antibodies for 6 hr with (w) or without (w/out) CECs at 1:1 or 1:2 ratios. (**C**) Representative plots, and (**D**) cumulative data of IFN-γand TNF-αexpression in CD4 and CD8 T cells following stimulation of PBMCs with S and N peptides of SARS-CoV-2 (2 μg/ml) for 6 hr with (w) or without (w/out) CECs at the indicated ratio. (**E**) Representative plots of CD107a in stimulated CD8 T cells with S and N peptides of SARS-CoV-2 for 6 hr in the absence or presence of CECs (1:1 ratio). (**F**) Representative plot, and (**G**) cumulative data showing proliferation of CD3 T cells as measured by CFSE dilution without (w/o) or with (w) CECs at 1:1 ratio following stimulation with S and N peptides of SARS-CoV-2 (2 μg/ml) for 3 days. (**H**) The correlation of total T cells, (**I**) CD4 T cells, and (**J**) CD8 T cells with the % CECs in the PBMCs of COVID-19 patients. Each point represents data from a patient.

These observations demonstrated the immunosuppressive properties of CECs in COVID-19 patients, potentially resulting in the impairment of both T and B cell effector functions.

### Progenitor CECs express SARS-Cov-2 receptor, ACE2

We have recently reported that HIV can both reside in CECs and that CECs can trans-infect CD4^+^ T cells^27^. Therefore, we speculated this might occur for SARS-CoV-2 virus. First, we examined whether CECs expressed the entry receptor for SARS-CoV-2, ACE2. We analysed ACE2 expression on CECs compared with immune cell subsets, and found that CECs were the dominant cells expressing ACE2 on their surface, followed by monocytes (Fig. 3A). Similar observations were made by Image stream analysis (Fig. 3B), and co-localization of ACE2 with CD71 and/or CD235a was noted (Fig. 3C). These observations were further re-confirmed by western blotting. The expression of ACE2 in different tissues/organs in mice was first confirmed (Extended Data Fig. 1I). Then, we examined CECs isolated from the peripheral blood of COVID-19 patients (purity of >95%, Extended Data Fig. 1J) and CECs from the placental tissues of humans for the presence of ACE2 (Fig. 3D). Next, we identified ACE2 expressing CECs as erythroid progenitors that express CD45 (Fig. 4A–C), using the gating strategy in Extended Data Fig.2A. The receptor-like tyrosine phosphatase CD45 is expressed on all nucleated hematopoietic cells including erythroid progenitors, and is downregulated when erythroid progenitors become mature RBCs^35^. CD45+CECs appeared to be the major ACE2 expressing cells in the peripheral blood of COVID-19 patients while other immune cells express negligible level of ACE2, except monocytes (Fig. 3A and 4C). Importantly, the expression and intensity of ACE2 was also significantly elevated on CD45+CECs compared to CD45-CECs and other immune cell lineages (Fig. 4D and 4E). In particular, the intensity of ACE2 was substantially higher in CD45hiCECs compared to their counterparts with lower CD45 expression (Fig. 4F and 4G). Nevertheless, the percentage of ACE2 expressing CECs varied during the course of disease (Extended Data Fig. 2B). In the absence of pathological conditions, erythrocytes are constantly produced under a highly orchestrated process regulated by multiple factors in the bone marrow and only enter the blood circulation once matured ^36^.

**Fig 3.**
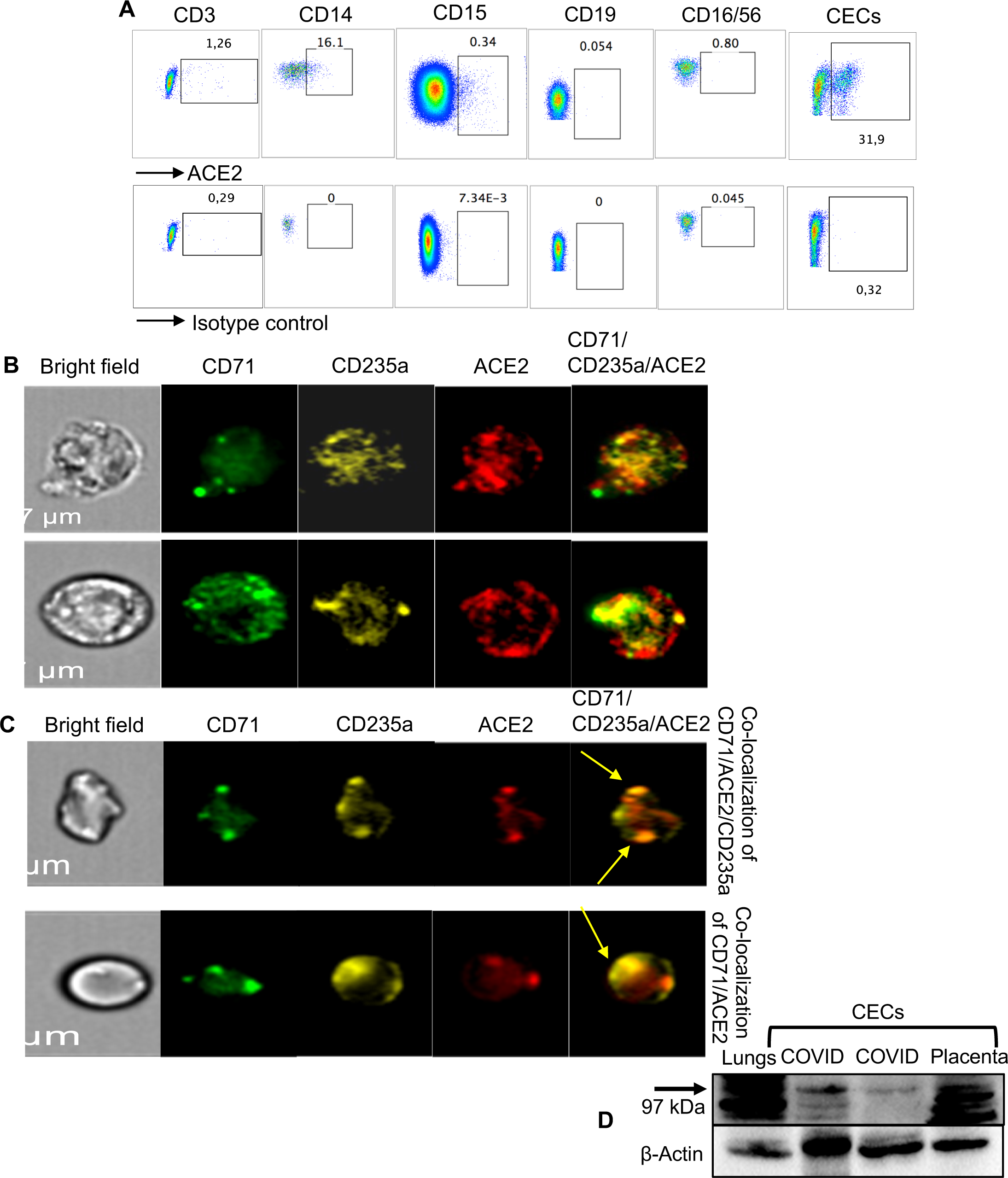
CECs possess surface expression of ACE2. (**A**) Plots showing the expression of ACE2 on CD3 T cells, CD14 monocytes, CD15 neutrophils, CD19 B cells, CD16CD56 NK cells and CECs compared to the isotype control antibody. (**B**) Image stream plots showing the expression of ACE2 on CECs, and (**C**) the co-localization of ACE2 with CD71/CD235a on CECs. (**D**) Western blot data showing the presence of ACEs protein in mouse lung tissue, CECs from two COVID-19 patients and CECs (~1×10^6^ cells) isolated from a placenta.

**Fig 4.**
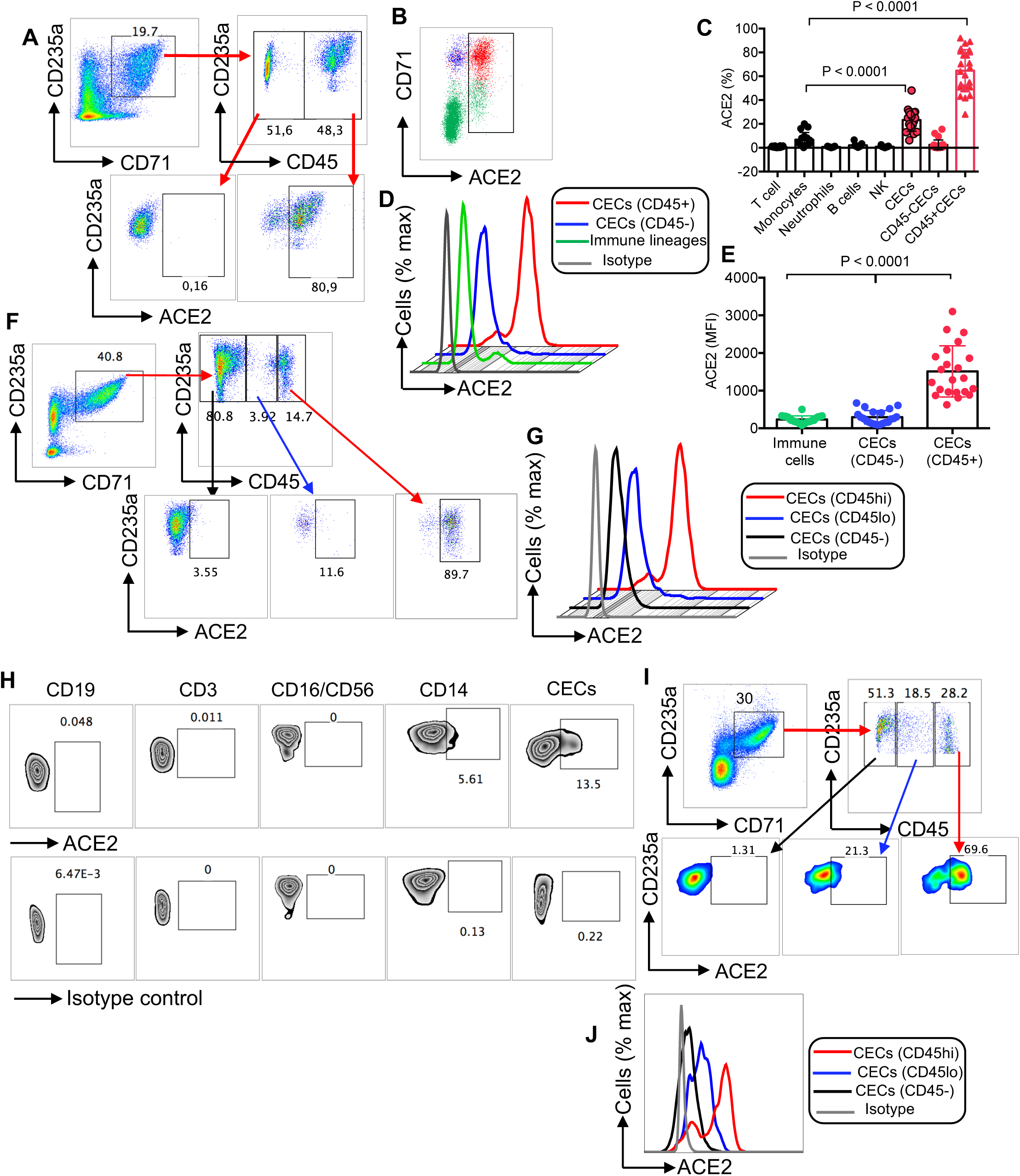
CD45+CECs from the peripheral blood of COVID-19 patients and human bone marrow exhibit the highest intensity of ACE2 expression compared to other immune cells. (**A**) Plots showing the expression of ACE2 on CD45-CECs vs. CD45+CECs of a COVID-19 patient. (**B**) Percent ACE2 surface expression on total CECs/immune cells. (**C**) Cumulative data of % ACE2 expressing cells among different immune cell lineages and CECs. (**D**) Representative plot, and (**E**) cumulative data of the intensity of ACE2 expression in CD45+CECs, CD45-CECs and immune cell lineages measured by mean fluorescence intensity (MFI). (**F**) Representative plots of % ACEs expression in CD45-, CD45lo and CD45hiCECs. (**G**) Plot showing the intensity of ACE2 expression in CD45hi, CD45lo and CD45-CECs compared to isotype control. (**H**) Plots showing the expression of ACE2 on CD19 B cells, CD3 T cells, CD16CD56 NK cells, CD14 monocytes and CECs compared to the isotype control antibody in the bone marrow aspirates of a healthy adult individual. (**I**) Representative plots of % ACEs expression in CD45-, CD45lo and CD45hiCECs of the bone marrow. (**J**) Histogram plot of the intensity of ACE2 expression in CD45hi, CD45lo and CD45-CECs of the bone marrow compared to isotype control. Each point represents data from a patient. Bar, mean ± one standard error.

We therefore decided to quantify the expression level of CD45 and ACE2 in CECs from the human bone-marrow. Since we were unable to obtain bone marrow aspirates from COVID-19 patients, as proof of concept, we examined bone marrow aspirates from non-COVID-19 patients for CD45 and ACE2 expression. We found that CECs were the most dominant ACE2 expressing cell type followed by monocytes (Fig. 4H and 4I). Similar to the peripheral blood CECs, the bone marrow CD45+hiCECs had the highest intensity of ACE2 (Fig. 4I and 4J). In addition to ACE2, we examined CD45+CECs for a second putative receptor for SARS-CoV2, CD147. We found that CD45+CECs possess the highest levels of CD147 expression when compared to other immune cells from COVID-19 patients (Extended Data Fig. 2C and 2D). Lastly, we found a higher expression of CD26 on CECs compared to RBCs (Extended Data Fig. 2E and 2F) and CD45+CECs had higher surface CD26 expression compared to CD45-CECs (Extended Data Fig. 2G). Since SARS-CoV-2 infection might lead to damage and lysis of these cells, we also measured soluble ACE2 in the plasma of COVID-19 patients. Interestingly, we found significantly higher levels of plasma ACE2 in patients with more moderate/severe disease than those with mild disease (Extended Data Fig.3A). As the disease progressed, plasma ACE2 levels also increased (Extended Data Fig. 3B). Longitudinal analysis in ICU patients also indicated a gradual increase in the plasma ACE2 with clinical progression (Extended Data Fig. 3C). Overall, these observations showed that CD45+CECs were the dominant ACE2/CD147 expressing cells, they also express CD26 in the peripheral blood of COVID-19 patients and soluble ACE2 was elevated in COVID-19 patients with moderate/severe disease.

### CD45+CECs also express SARS-CoV-2 co-receptor, TMPRSS2

Recent evidence indicates that viral entry into target cells depends not only on the binding of the spike (S) protein to ACE2 but also requires S protein priming by the cellular serine protease TMPRSS2^6^. Thus, we found that CD45+CECs both express TMPRSS2 (Fig. 5A and 5B and Extended Data Fig. 3D) and co-express ACE2 and TMPRSS2 (Fig. 5A). These observations were confirmed by image stream analysis (Fig. 5C and Extended Data Fig. 3E and 3F). Moreover, the intensity of TMPRSS2 was significantly greater on CD45+CECs compared to CD45-CECs and other immune cell lineages (Fig. 5D and 5E). Of note, a small subset of the immune cell lineages, mainly CD14 monocytes also express TMPRSS2 (Fig. 5F and Extended Data Fig. 3D). These observations were further re-confirmed by western blotting. Initially, we confirmed the expression of full length (54 kDa) and the cleavage fragment (25 kDa) of TMPRSS2 in mice tissues/organs compared to the positive control (human colorectal adenocarcinoma grade II cell line) (Extended Data Fig.4A), however, the cleaved product appeared to be smaller than 25 kDa in mice tissues. Western blot confirmed the expression of TMPRSS2 in CECs isolated from COVID-19 patients and human placental tissues (Fig. 5G). Interestingly, when the same number of cell lysate was loaded, TMPRSS2 protein level was higher in CECs compared to immune cell linages (CECs-) (Extended Data Fig. 4B). Of note, β-actin level appeared to be lower in CECs compared to other immune cells. Similarly, bone marrow CECs also possess the surface expression of TMPRSS2 (Extended Data Fig. 4C). Taken together, these results confirm the presence/surface expression of TMPRSS2 and its co-expression with ACE2 in CECs.

**Fig 5.**
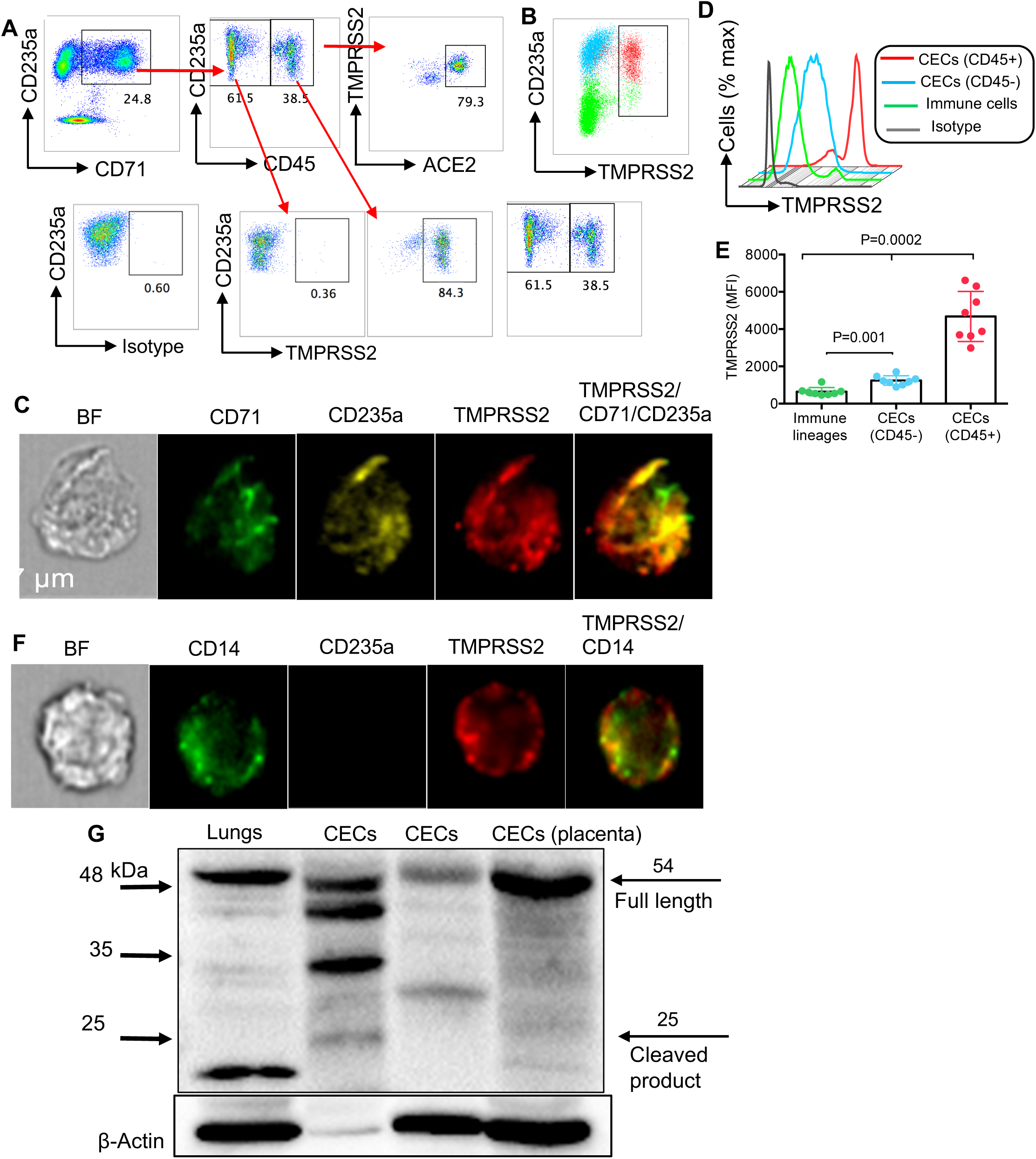
CD45+CECs from the peripheral blood of COVID-19 patients are the dominant TMPRSS2 expressing cells. (**A**) Plots showing the expression of ACE2 on CD45-CECs vs. CD45+CECs and the co-expression of ACE2/TMPRSS2 on CD45+CECs of a COVID-19 patient. (**B**) Percent TMPRSS2 surface expression on total CECs/immune cells. (**C**) Image stream plots of TMPRSS2 expression on CECs (**D**) Histogram plot, and (**E**) cumulative data of the intensity of TMPRSS2 expression in CD45+CECs, CD45-CECs and immune cell lineages measured by mean fluorescence intensity (MFI). (**F**) Image stream plots of TMPRSS2 expression on CD14 monocytes of a COVID-19 patient. (**G**) Western blot data showing the presence of TMPRSS2 protein in mouse lungs tissue, CECs from two COVID-19 patients and CECs (~1×10^6^ cells) isolated from a placenta tissue. Each point represents data from a patient. Bar, mean ± one standard error.

### CECs get infected with SARS-CoV-2 and dexamethasone reduces their infectivity *in vitro*

In light of ACE2 and TMPRSS2 expression/co-expression and CD147/CD26 expression on CECs, we hypothesized that these cells could get infected by SARS-CoV-2. To test this hypothesis, we first examined the interaction of CECs with the spike receptor binding domain. We found that the spike binding domain (conjugated by a fluorescent dye) bound ACE2 on CD45+CECs whereas it did not bind CD45-CECs (Fig. 6A). This was further confirmed by Image stream analysis (Fig. 6B). The ability of CECs from COVID-19 patients to be infected by SARS-CoV-2 using the traditional approach versus magnetofection was tested as we have reported for HIV-1^37,38^. We were able to detect viral RNA produced in cell culture supernatants as well as viral RNA in the cells 24 hrs after infection using both methods, but the magnetofection was more efficient (Fig. 6C and 6D). While viral RNA levels were not high compared to VeroE6 cells, they were > 2 logs higher than the final wash after infection and infectious SARS-CoV2 was produced by these cells. Since it has been recently shown that monocytes can be infected with SARS-CoV2^39^, we compared the amount of infection in CECs to that in monocytes. We found that the amount of viral RNA in CECs and produced by CECs was similar to monocytes (Fig. 6E and 6F). To examine the infectivity of CEC’s obtained from a SARS-CoV2 naïve source we infected CECs isolated from human placenta with SARS-CoV2, because placenta is physiologically enriched with CECs^33–34,40,41^ possessing ACE2 and TMPRSS2 (Fig. 3D, Fig. 5G). Similar to CECs from COVID-19 patients, placental CD45^+^CECs had substantial surface expression of ACE2 (Extended Data Fig.5A and 5B) and TMPRSS2 (Extended Data Fig. 5C). We found that similar to CEC’s from COVID-19 patients, CEC’s from placenta contained viral RNA after infection and secreted viral RNA into the cell culture supernatants (Fig. 6G and 6H). These results indicate that CEC’s can be directly infected by SARS-CoV2.

**Fig 6.**
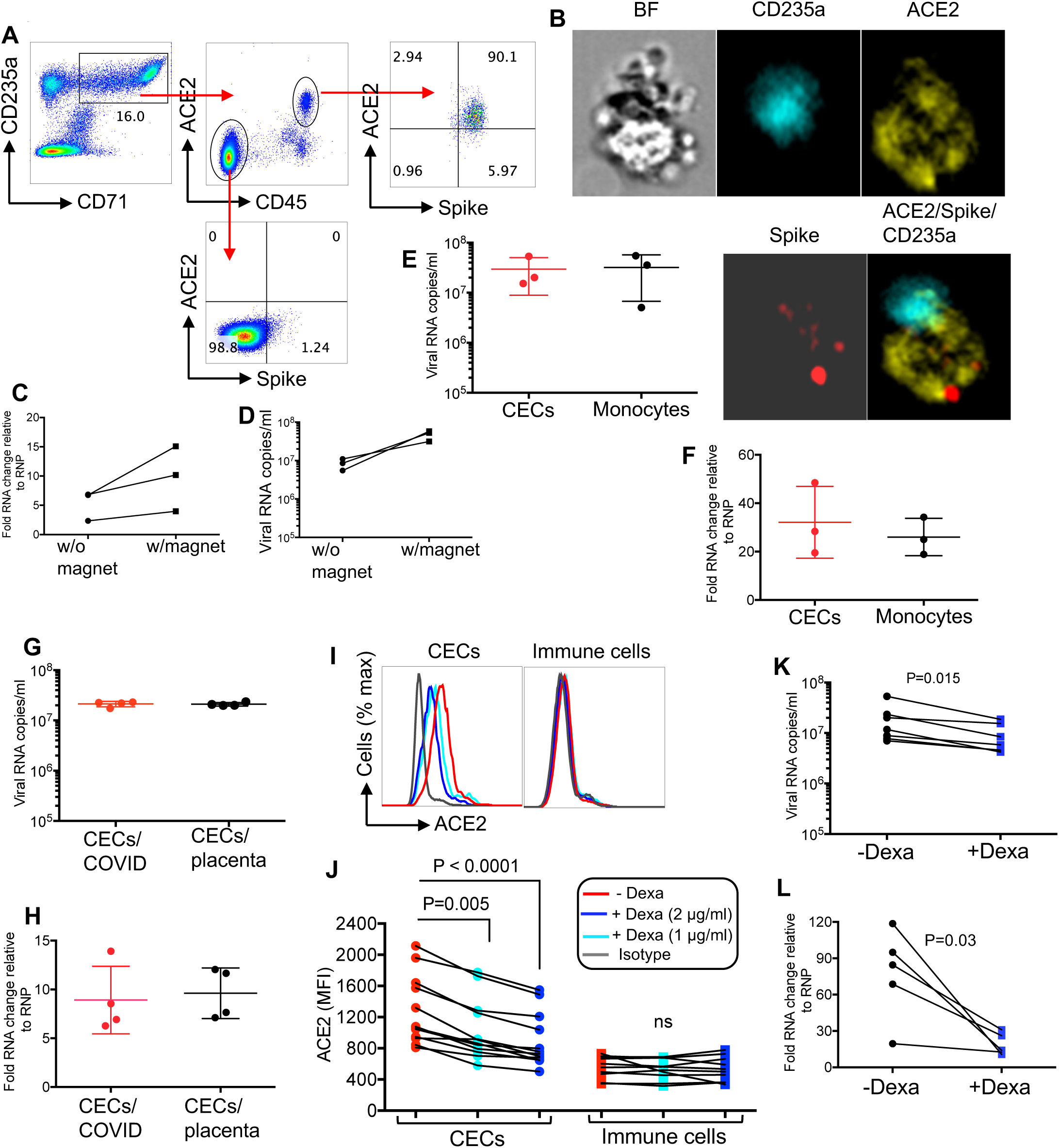
CECs are permissible to SARS-CoV-2 infection which can be reversed in part by dexamethasone. (**A**) Plots showing co-expression of spike binding domain with ACE2 on CD45+CECs but not CD45-CECs. (**B**) Image stream plots of spike protein interaction/binding with ACE2 on CECs. (**C**) Cellular RNA level in CECs, and (**D**) viral RNA copies in culture supernatants of CECs without (w/o) or with (w) magnetofection with SARS-CoV-2. (**E**) Viral RNA copies in the culture supernatant of total CECs or monocytes of COVID-19 patients infected with SARS-CoV-2 and measured by qPCR 24 post infection. (**F**) Cellular RNA changes relative to the housekeeping gene (RNP) in cell pellet of CECs and monocytes of COVID-19 patients infected with SARS-CoV-2 and measured by qPCR 24 post infection. (**G**) Viral RNA copies in culture supernatants of total CECs of COVID-19 patients versus CECs of the placental tissues of healthy deliveries 24 hr post infection with SARS-CoV-2. (**H**) Cellular fold RNA changes relative to the housekeeping gene (RNP) in cell pellet of total CECs from COVID-19 patients or the placental tissues of healthy deliveries 24 hr post infection with SARS-CoV-2. (**I**) Histogram plots, and (**J**) cumulative data of the expression of ACE2 on total CECs and immune cells lineages of COVID-19 patients treated with or without dexamethasone (1 or 2 μg/ml) overnight. (**K**) Viral RNA copies in culture supernatants of CECs either untreated or treated with dexamethasone (2 μg/ml) for 24 hr before infection with SARS-CoV-2 as measured 24 post infection. (**L**) Cellular fold RNA changes relative to the housekeeping gene (RNP) in cell pellet of CECs either untreated or treated with dexamethasone (2 μg/ml) for 24 hr before infection with SARS-CoV-2 as measured 24 hr post infection. Each point represents data from cells of a patient, infection studies are representative of three independent experiments. Bar, mean ± one standard error.

Erythroid progenitors possess a glucocorticoid receptor which enhances the response to erythropoietin (Epo) ^42^. Glucocorticoids such as dexamethasone are known to aid in the treatment of Epo-resistant anemia by stimulating self-renewal of progenitors ^43,44^. Since COVID-19 patients and primate models of SARS-CoV-2 infection are generally anemic^45,46^ and little is known about the mechanism associated with the therapeutic effects of dexamethasone in severely ill COVID-19 patients^47^; we reasoned that the enhanced maturation of expanded CECs in severe disease might be one molecular mechanism for the lower mortality rate in patients receiving dexamethasone. We first examined the effect of dexamethasone in mice. Since young mice possess a higher frequency of CECs in their spleens compared with adults^30^, we treated young mice (15 days) with dexamethasone (1μg/g body weight by i.p.) and 2 days later collected their spleens for the quantification of CECs. A significant reduction in the frequency of CECs in treated versus control animals was observed (Extended Data Fig. 5D and 5E), suggesting that dexamethasone enhances the maturation of CECs to mature RBCs. We then examined whether CECs obtained from the peripheral blood of COVID-19 patients, displayed the same phenomena. We treated total PBMCs and/or isolated CECs with 1 and 2 μg/ml dexamethasone overnight^48^. We observed that such treatment enhanced the maturation of CECs which resulted in the downregulation of ACE2 in a dose-dependent manner (Fig. 6I and 6J), but did not affect the ACE2 expression on other immune cells lineages (Fig. 6I and 6J), indicating that dexamethasone specifically modulates ACE2 expression in CECs; possibly due to the presence of glucocorticoid receptors on CECs. A similar effect was observed for TMPRSS2 expression on CECs following treatment with dexamethasone (Extended Data Fig. 5F). In parallel we examined whether atorvastatin, due to its immunomodulatory properties as we previously reported^38,49^, can modulate ACE2 expression. However, this was not the case (Extended Data Fig. 5G). Since dexamethasone downregulated the expression of ACE2, we reasoned it might also reduce the permissibility of CECs to SARS-CoV-2 infection. When we pre-treated CECs with 2 μg/ml dexamethasone for 24 hrs prior to SARS-CoV2 infection, there was a significant reduction in the viral RNA in cell culture supernatant and intracellular viral RNA compared to untreated cells (Fig. 6K and 6L). Importantly, this was not the case for monocytes when treated with dexamethasone (Extended Data Fig. 5H and 5I). Taken together, these observations indicate that CECs can be infected by SARS-CoV-2, and this can be partially inhibited by dexamethasone.

## Discussion

The present study provides the first comprehensive analysis of the role of erythroid progenitors/precursors (CECs) in COVID-19 infection. We identified significant expansion of CECs in the peripheral blood of COVID-19 patients, in particular, in those with moderate or severe disease. Interestingly, we found that the expansion of CECs was associated with disease severity and ICU admission. CECs have a wide range of immunomodulatory properties and via cell: cell interactions or soluble mediators such as TGF-β, ROS and arginase II, and can suppress effector T cell and B cell functions ^23–25,27,29,34^. Given the association of CECs with disease severity, further analyses were performed. Consistent with previous reports in the context of infection (e.g. HIV) and cancer^26–29,31^, CECs from COVID-19 patients exhibited substantial expression of arginase II and ROS. In addition, arginase I expression has not previously been reported in CECs. These capabilities not only enabled CECs to exert a global immunosuppression effects on T cells but they also significantly impaired cytokine production, proliferation and degranulation capacities of antigen-specific T cells *in vitro*. These observations were further confirmed *in vivo* by the presence of a negative correlation between the frequency of T cells with the percentages of CECs in the peripheral blood of COVID-19 patients. Therefore, the pathological abundance of CECs in COVID-19 patients may in part explain the mechanism underlying a wide range of changes in the frequency and functionality of different immune cells in these patients ^4,13^. One striking observation was that CECs from COVID-19 patients appear to have a different membrane structure. For example, measuring arginase-II activity in human CECs has been technically impossible as CECs are lysed when exposed to the fixation/permeabilization buffer required for intracellular staining. Surprisingly, this was not the case for CECs isolated from COVID-19 patients and indeed, these CECs were resistant to permeabilization. We also observed increased number of light weight RBCs (CD235a+CD71-) in the circulation of these patients. This suggests that SARS-CoV-2 infection or the immunopathology associated with COVID-19 disease can modify RBCs and/or CEC structural components, and was illustrated by a recent report that COVID-19 infection is associated with RBC structural protein damage and modifications in RBC membrane lipids ^17^. Although this study was conducted on mature RBCs, it is possible that COVID-19 infection can also alter CEC deformability. As such, the abundance of these unwanted guests and the possible altered structural proteins following infection with SARS-CoV-2 may contribute to other serious complications (such as thromboembolic and coagulopathic) commonly observed in COVID-19 patients ^50^. The role of RBC morphology and deformability in clot formation has been widely studied ^51,52^. Although thrombosis is likely multifactorial in nature, involving vasculature, platelets and dysfunctional RBCs, it will be of interest to determine if expanded CECs alike their older siblings are involved within thrombi in COVID-19 patients. The severely low oxygen saturations observed in critically ill patients^53^; particularly without substantial damage to the lungs^54^ suggests that SARS-CoV-2 may affect oxygenation via paths unrelated to pulmonary function. Elevated cytokines (e.g. IL-1, IL-6 and TNF-α) may influence erythropoiesis^55^ and increase the permissibility of RBCs to oxidant stress-induced lysis in COVID-19 patients^17^. In light of the above, there is the possibility that SARS-CoV-2 infection directly or indirectly invades RBCs in the periphery or the bone marrow resulting in their enhanced lysis and anemia^45–46,56^. This can result in stress erythropoiesis and subsequently abundant CECs in the periphery due to passive incontinence of hematopoietic cells from the bone marrow^25,57^. Although the lack of organelles in RBCs precludes viral survival/replication, this might not be true for CECs. Recently, we reported that HIV can reside and possibly replicate within CECs^31^. In support of this hypothesis, we identified CECs as the major ACE2 and TMPRSS2 expressing cells followed by monocytes. Other immune cells in the peripheral blood of COVID-19 patients had negligible expression of both SARS-CoV-2 entry receptors. Further, the receptor-like tyrosine phosphatase CD45, is expressed on all nucleated hematopoietic cells including erythroid progenitors^35^ and gets downregulated as erythroid progenitors mature into RBCs^35^. Of note, CD45+CECs were the dominant ACE2/TMPRSS2 expressing cells compared to their CD45-CEC counterparts. In this setting, we speculated that erythroid progenitors in the bone marrow should express ACE2 and TMPRSS2. Indeed, this was the case and once again we observed CD45+CECs as the dominant cells in terms of ACE2 and TMPRSS2 expression in the human bone marrow. In addition, we found that CECs from COVID-19 patients, in particular, CD45+CECs, possess the highest intensity of CD147/CD26 expression, another SARS-CoV-2 receptor^18^. Taken together, these observations suggest that CECs could be an attractive target for SARS-CoV-2. It is worth noting, that soluble ACE2 was elevated in plasma of COVID-19 patients and it was more prominent in patients with severe disease. It is possible to speculate that this soluble ACE2 may shed from CECs following lysis, however, further investigations are needed to identify the source and the role of soluble ACE2 in these patients. More importantly, CECs, in particular their CD45+ progenitors possess nuclei and other organelles that can support viral survival and replication. Therefore, we first confirmed the binding of SARS-CoV-2 spike with ACE2 on the surface of CD45+CECs. These observations led us to determine if CECs can get infected with SARS-CoV-2 and more importantly whether the virus replicates in these cells. We found that the infectivity of CECs to SARS-CoV-2 was comparable to monocytes. Despite the similar level of viral infection in these two cell subsets, we believe CECs are more permissible to infection compared to monocytes for multiple reasons. 1) CECs are the most abundant cells in the peripheral blood of COVID-19 patients especially those with a moderate/severe disease. 2) CECs contain the highest ACE2, CD147 and TMPRSS2 expression. 3) Given a lower proportion of nucleated CECs but similar viral RNA copies were observed in the culture supernatant and cell pellet of CECs and monocytes, this suggests more efficient viral infectivity/replication in CECs. The CD45+CECs population comprises approximately 20-80% of CECs while monocytes are 100% nucleated and can support viral replication. Thus, the infectivity of RBC progenitors to SARS-CoV-2 infection may explain one potential mechanism for the observed hypoxia in COVID-19 patients. As such, higher percentages of CECs in these severely ill patients is indicative of stress hematopoiesis. We hypothesize that this phenomenon might be due to the elimination of infected/damaged CECs by lysis or/and phagocytosis. However, further studies are required to confirm these observations by detecting viral proteins or infective viral particles in CECs of COVID-19 patients.

On the other hand, the immunosuppressive properties of CECs may be beneficial to COVID-19 patients since hyper-inflammation and cytokine storm is associated with disease severity^13^. As such, CECs might appear protective at the early stage of disease to prevent a robust innate immune response. Nevertheless, expansion of CECs in the peripheral blood of COVID-19 patients coincides with the disease progression, which is the time for the induction of an efficient adaptive immune response. Therefore, the absence of CECs at the early stage of disease deprives the host from their highly desired immunosuppressive properties but instead their appearance later can compromise T cell effector functions and antibody production. The uncontrolled inflammatory response can itself damage the lungs via the excessive release of proteases, reactive oxygen species, and pro-inflammatory cytokines^58^. In agreement with this concept, several immunosuppressive strategies are recommended for the treatment of COVID-19 patients. Although the administration of systemic corticosteroids for COVID-19 patients was initially not supported by the WHO guidelines^13^, several trials are under way for the efficacy of such treatment options^58^. In fact, a recent randomized clinical trial has shown that dexamethasone reduces deaths by one-third in patients on ventilators and by one-fifth in those receiving oxygen without invasive ventilation ^47^. In addition to the anti-inflammatory function of dexamethasone, it influences haematopoiesis and promotes the maturation of erythroid cells ^59^. This may be supported by our observation that dexamethasone reduced ACE2/TMPRSS2 expression by enhanced CECs maturation. In this respect, dexamethasone-mediated downregulation of ACE2/TMPRSS2 may explain the reduced susceptibility of CECs to SARS-CoV-2 infection. Knowledge gained from this study may illuminate the pivotal role CECs play in COVID-19 pathogenesis. In addition, this study provides mechanistic rationale for the clinical use of dexamethasone in COVID-19 patients, in particular, in those with severe disease. It is therefore, tempting to speculate that immunosuppressive drugs might be harmful when given in the induction phase of immune response. However, considering the massive expansion of CECs in COVID-19 patients, targeting these cells with medications such as dexamethasone might be beneficial rather than detrimental for the patient. It appears that dexamethasone may not only attenuate the hyperactive immune response but also protects CECs from the virus, and enhancing their maturation and preventing hypoxia.

Given the high surface expression of ACE2 and TMPRSS2 on placental CECs and their permissibility to ARS-CoV-2 infection, raises the possibility of vertical viral transmission seems possible. Also, CECs become abundant in the peripheral blood of pregnant women at the later stage of pregnancy^33–34,40^, which in part may support the reported viremia and placental transmission of SARS-CoV-2^60^. This was challenged due to negligible co-expression of ACE2 and TMPRSS2 by placental cell types^61^. However, mRNA expression does not necessarily correlate with the protein expression pattern. More importantly, the expression of ACE2 and TMPRSS2 on CECs as the major cells in placenta was not studied in this report. Thus, the abundance of CECs expressing ACE2 and TMPRSS2 in the peripheral blood of pregnant mothers and placenta tissue supports transplacental transmission of SARS-CoV-2 infection. We are aware of the study limitations such as low sample size, especially in infected patients who do not require hospitalization. Moreover, due to blood sample size we were restricted in terms of the number and depth of analyses. For example, we would have liked to perform additional infection assays (e.g. drug titration, time points, blood group). Moreover, we were unable to perform longitudinal analyses of CECs functions/infectivity overtime. A larger sample would also be required to evaluate whether dexamethasone treatment reduces the viral load in patients (e.g. lungs) and if it influences the expression of ACE2 in different tissues.

## Methods

### Human sample collection and processingn

Blood samples were collected from hospitalized COVID-19 patients in different hospitals in Edmonton, Alberta. All COVID-19 patients were SARS-CoV-2 positive by qRT-PCR assay specific for viral RNA-dependent RNA polymerase and envelope transcripts detected using a nasopharyngeal swab. The Human Research Ethics Board (HREB) at the University of Alberta approved the study (Pro00099502). Waiver of consent was obtained by the HREB for those patients admitted to the ICU but a verbal consent was required from all other patients. Wet consent was not required due to logistics and the possible risk of viral transmission. Fresh peripheral blood mononuclear cells (PBMCs) were isolated over Ficoll-Hypaque gradients. For CECs or mock depletion samples were stained using anti-CD71 or isotype control biotin-conjugated antibody and fractioned using streptavidin linked magnetic beads (Miltenyi Biotec) according to our previous reports ^24–26,31,34^. Normally, the isolated CECs have a purity > 95% (Extended Data Fig. 1E and 1J).

### Antibodies and flow cytometry

Fluorophore or biotin-conjugated antibodies with specificity to human cell surface antigens and cytokines were purchased mainly from BD Biosciences or Thermo Fisher Scientific and in some occasions from other suppliers as indicated below. Specifically, the following antibodies were used: anti-CD3 (HIT3a), anti-CD4 (RPA-T4), anti-CD8 (RPA-T8), anti-CD45 (H-130 or 2D1), anti-VISTA (B7H5DS8), anti-107a (H4A3), anti-PD-L1 (MIH1), anti-CD147 (8D12), anti-CD16 (B73.1), anti-CD56 (B159), anti-CD15 (HI98), anti-CD14 (M5E2), anti-IL-2 (MQ1-17H12), anti-TNF-α(MAB11), anti-IFN-γ(4S.B3), anti-CD71 (MA712), anti-CD235A (HIR2) and arginase I (IC5868N/R&D). ROS (Sigma) and arginase II (abcam) staining were performed per the manufacturer’s protocols and our previous reports^24–31,34^. In addition, anti-ACE2 (535919) from R&D, and anti-TMPRSS2 (EPR3862) from abcam were used for staining. The SARS-CoV-19 spike receptor binding domain protein was purchased from VIROGEN (Cat#00224-V), conjugated with dye using Fluorescent protein labeling kit according to the manufacturing protocol (Thermo Fisher Scientific) for related studies. Besides, Live/dead fixable dead cell stains (ThermoFisher) were used to exclude dead cells in flow cytometry. Paraformaldehyde fixed cells were acquired by flow cytometry using a LSRFORTESSA flow cytometer (BD) and analyzed with FlowJo software.

### Co-culture and stimulation

For *in vitro* intracellular cytokine staining, PBMCs were cultured and stimulated with anti-CD3/CD28 in RPMI media supplemented with 10% FBS for 6 hours in the presence or absence of CECs according to our previous report ^27^. For co-culture, a fixed number (1 × 10^6^) of PBMCs were seeded into 96 well round bottom plates individually or together with autologous CECs at 1:1 ratio, Brefeldin A (10 μg/ml) was added at the same time. In other experiments, PBMCs were stimulated with SARS-CoV-2 peptide pools of S and N (2 μg/ml) (Miltenyi Biotec) in the presence or absence of CECs. Similar approach was used for proliferation assay, in brief, PBMCs were labelled with CFSE and stimulated with peptide pools in the presence or absence of CECs (1:1 ratio) for 3 days according to our previous protocols^34,62^.

### ELISA

IL-33 (Novus Biologicals) and ACE2 DuoSet ELISA (R&D) were performed on frozen plasma samples of patients and healthy controls according to the manufacturing protocol.

### Western blot analyses

Cells and tissues were lysed in lysis buffer supplemented with a protease inhibitor cocktail (Sigma-Aldrich) and protein concentration was determined using a BCA assay kit (Thermo Fisher Scientific). Protein samples were separated by electrophoresis on either 7%, 17% or 4-15% gradient polyacrylamide gels and then transferred to PVDF membranes. The membranes were blocked with 5% milk and incubated with anti-ACE2 (Abcam, ab15348), anti-TMPRSS2 (Abcam, ab242384) and anti-β-actin (Sigma, A2228) antibodies using dilutions 1:1,000. HT29 (human colorectal adenocarcinoma grade II) whole cell lysate was used as a positive control for TMPRSS2 (Abcam, ab3952). Full length predicated at 54 kDa and the cleavage fragment at 25. Next, membranes were incubated with the appropriate HRP-conjugated secondary antibodies and developed using an enhanced chemiluminescence detection kit (Thermo Fisher Scientific).

### Virus infection and quantification

SARS-CoV-2 stocks were pre-incubated with ViroMag transduction reagent (OZ Biosciences) as reported elsewhere^31–37,38^. To remove background virus, cells were washed and pelleted five times with 15 ml of media, a sample of the last wash was taken to measure remaining background viral RNA (~10^5^ viral copy). In some experiments, target cells were pre-treated with dexamethasone (2 μg/ml) overnight prior to the infection. Following a 24-hour incubation at 37°C, a sample of the culture supernatant was taken to measure extracellular virus production. RNA was extracted using QIAamp Viral RNA mini kit (Qiagen). To measure intracellular RNA, cells were first washed and pelleted five times with 15 ml of PBS then lysed with QIAzol reagent (Qiagen), RNA was extracted according to manufactures directions. Reverse transcription was carried out using Superscript IV Vilo master mix (Invitrogen). Quantitative PCR was carried out using primers and probe designed by the United States center for disease control and prevention: for the N gene (N2 primers) of SARS-CoV-2 and RNAse P housekeeping gene (IDT cat#10006606). A standard curve was generated using dilutions of positive control standards from CDC (IDT cat # 10006625).

### Statistical analysis

Statistical comparisons between various groups were performed by using t-test and Mann-Whitney tests (as appropriate) using PRISM, Graph Pad software. Also, differences were evaluated using One-Way ANOVA followed by Tukey’s test for multiple comparisons. Correlation analysis was performed using spearman test. Results are expressed as mean± SEM. P-value <0.05 was considered statistically significant.

## Acknowledgements

We thank our study volunteers for providing samples and supporting this work and the clinical staff for their dedication to this research. This work was supported mainly by a grant from FASTGRANT.com. We also acknowledge the Canadian Institutes of Health Research (CIHR) for the financial support of Elahi lab.

## Author contributions

S.S. performed most of the experiments and analyzed the data. L.X. performed some of the experiments and analyzed the data. M.O. who contributed in patients’ recruitment. W.S. as a clinician scientist (in critical care medicine and infectious disease) who identified and recruited patients for the study. J. S. performed viral infection experiments. M.J. provided assistant and guided viral infection studies. L.T. advised on infection experiments. O.O. performed all the western blotting studies. S.E. conceptualized, designed, secured funding and resources, analyzed the data, supervised all of the research and wrote the manuscript. All authors revised and edited the manuscript.

