## Supplemental Figures for "Erythroid precursors and progenitors suppress adaptive immunity and get invaded by SARS-CoV-2"

Extended Data Fig. 1

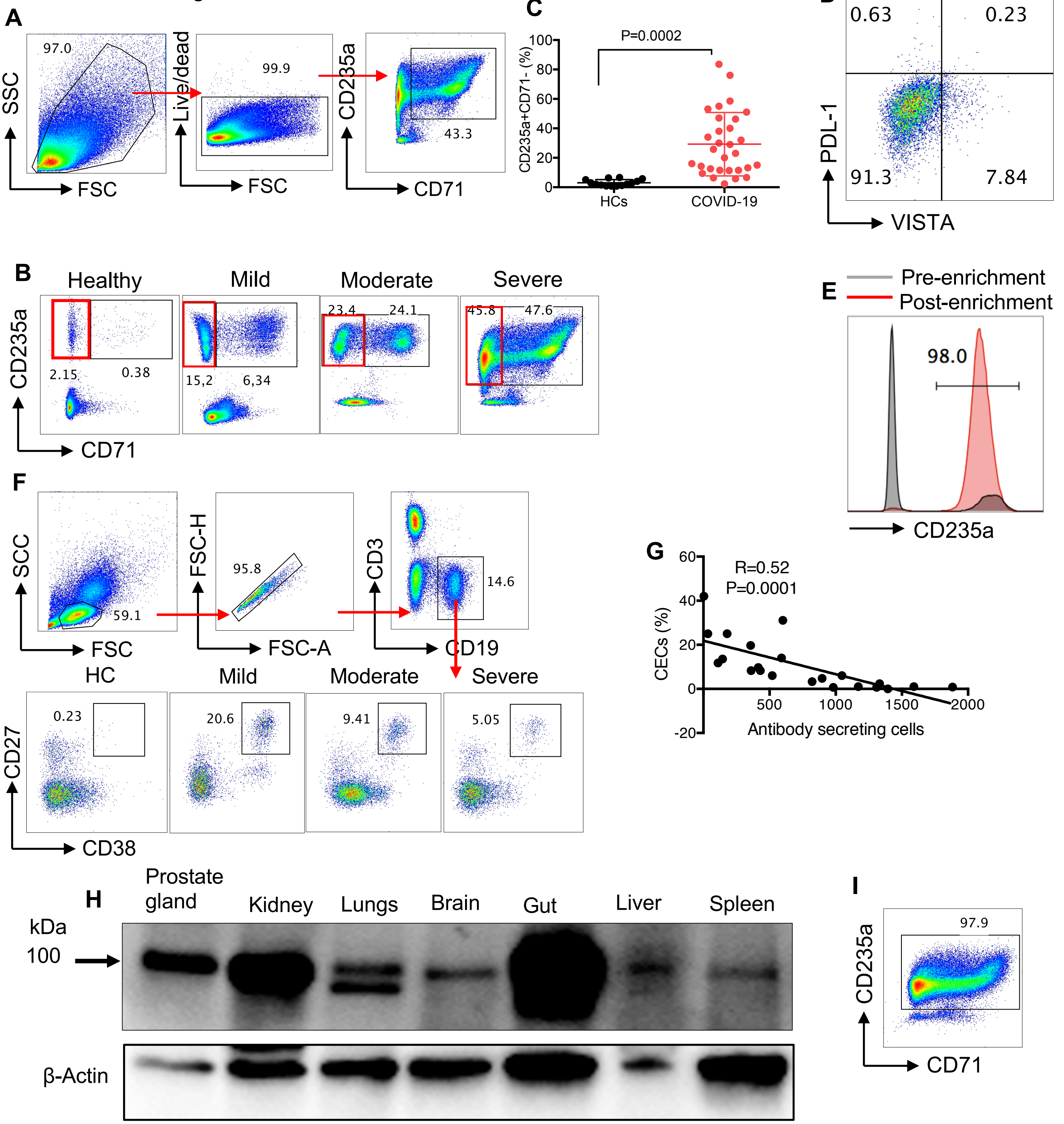

Extended Data Fig. 2

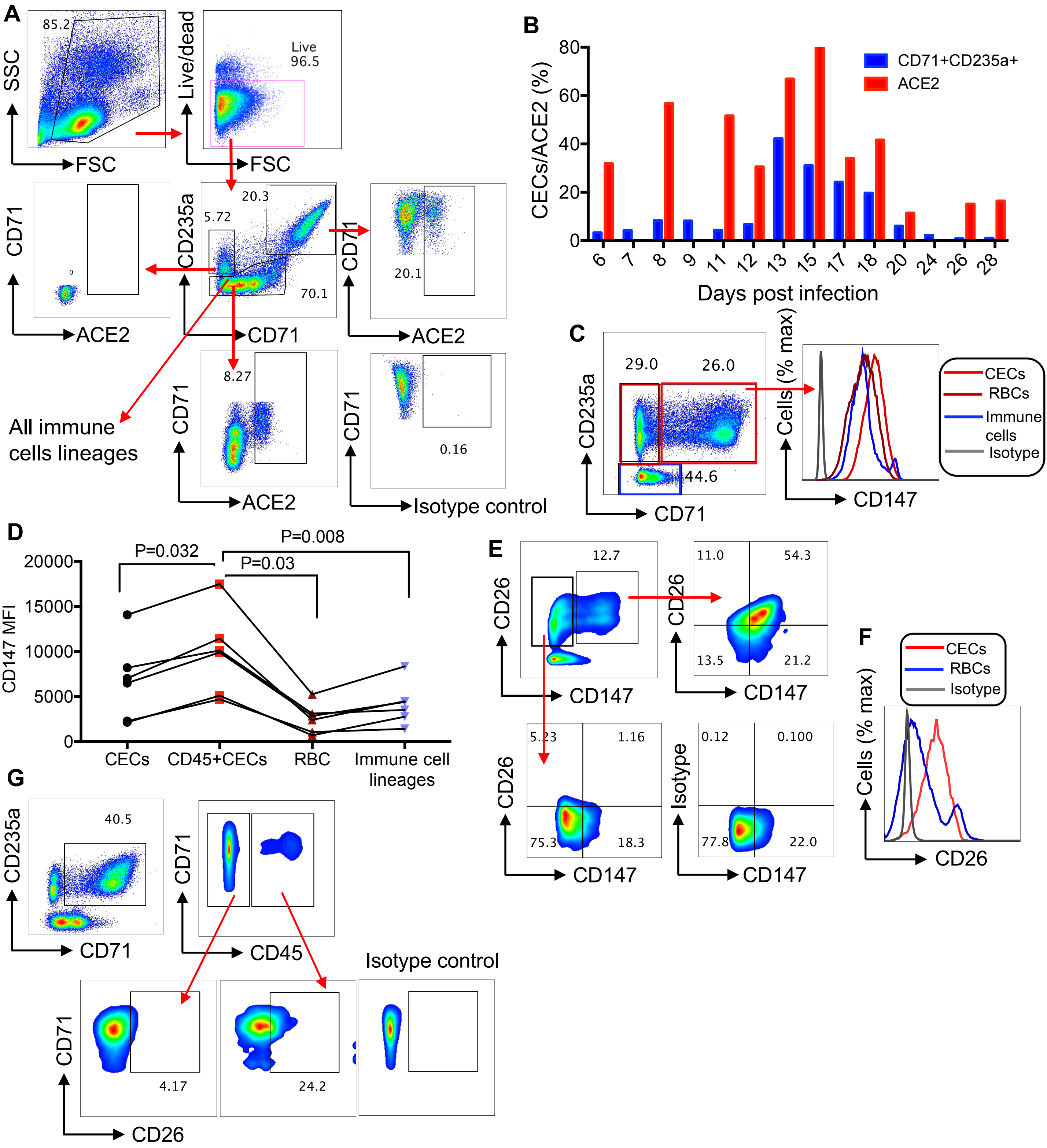

Extended Data Fig. 3

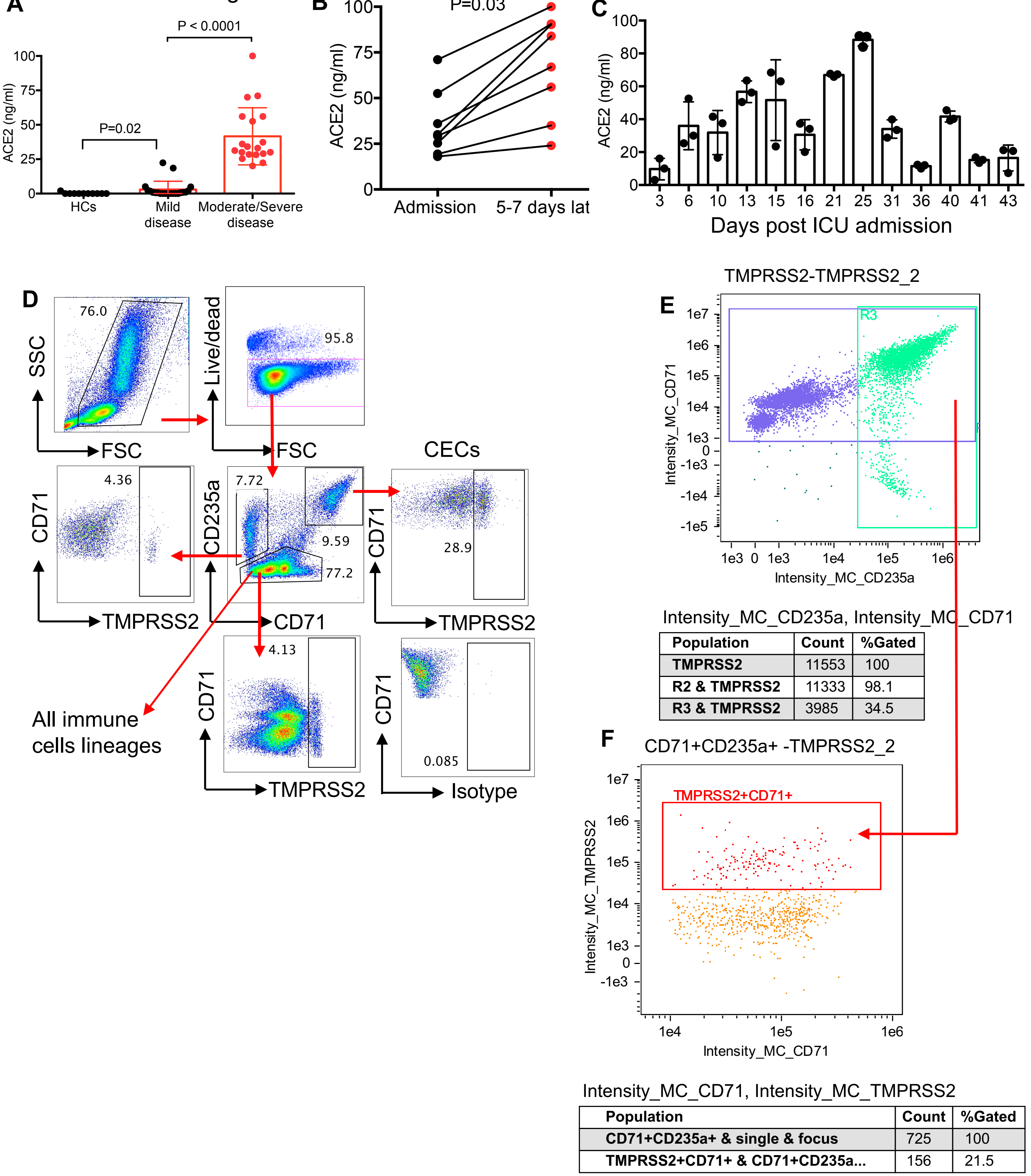


Intensity_MC_CD235a, Intensity_MC_CD71

| Population | Count | %Gated |
|---|---|---|
| TMPRSS2 | 11553 | 100 |
| R2 & TMPRSS2 | 11333 | 98.1 |
| R3 & TMPRSS2 | 3985 | 34.5 |

Intensity_MC_CD71, Intensity_MC_TMPRSS2

| Population | Count | %Gated |
|---|---|---|
| CD71+CD235a+ & single & focus | 725 | 100 |
| TMPRSS2+CD71+ & CD71+CD235a... | 156 | 21.5 |

Extended Data Fig. 4

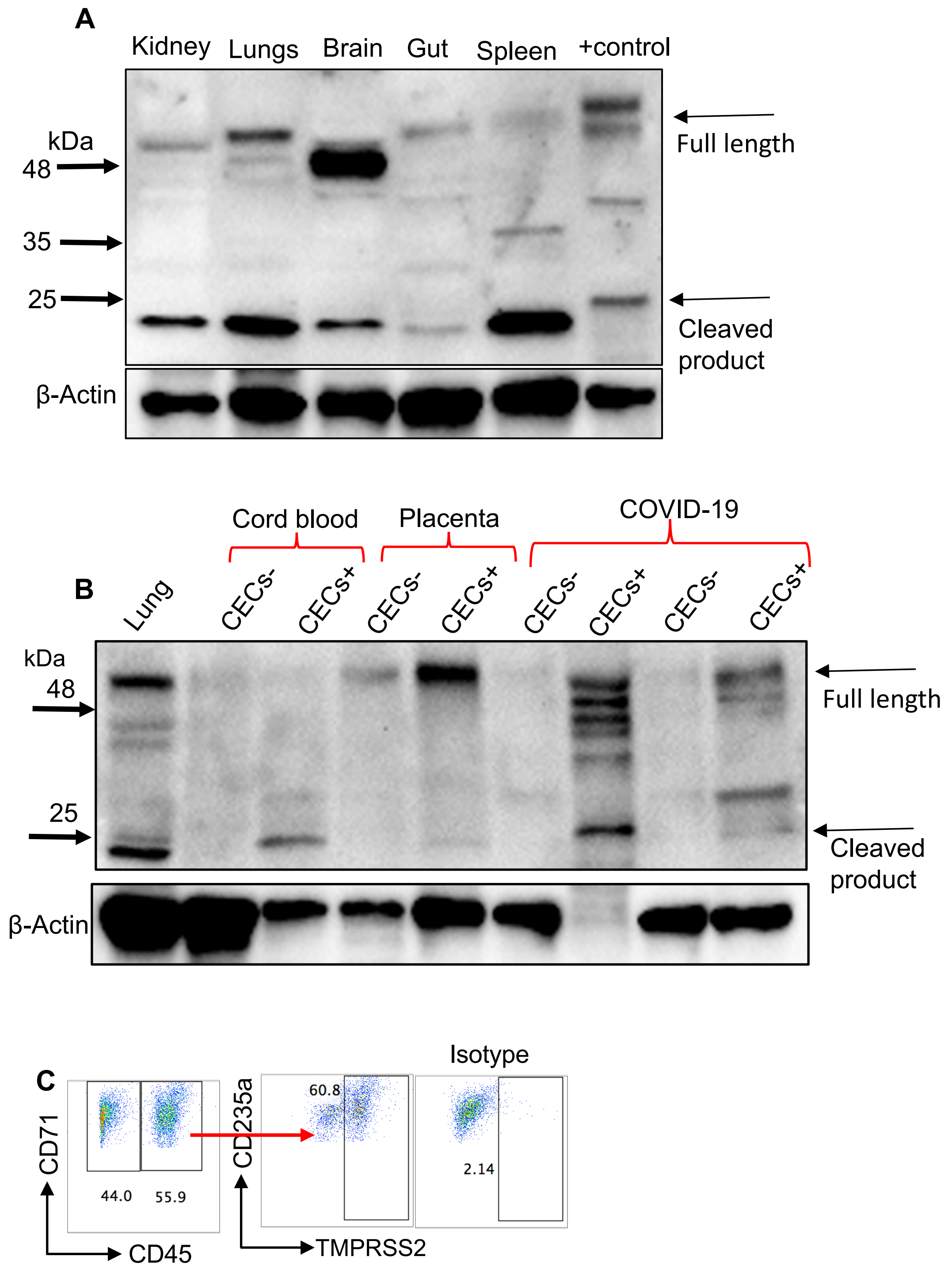

Extended Data Fig. 5

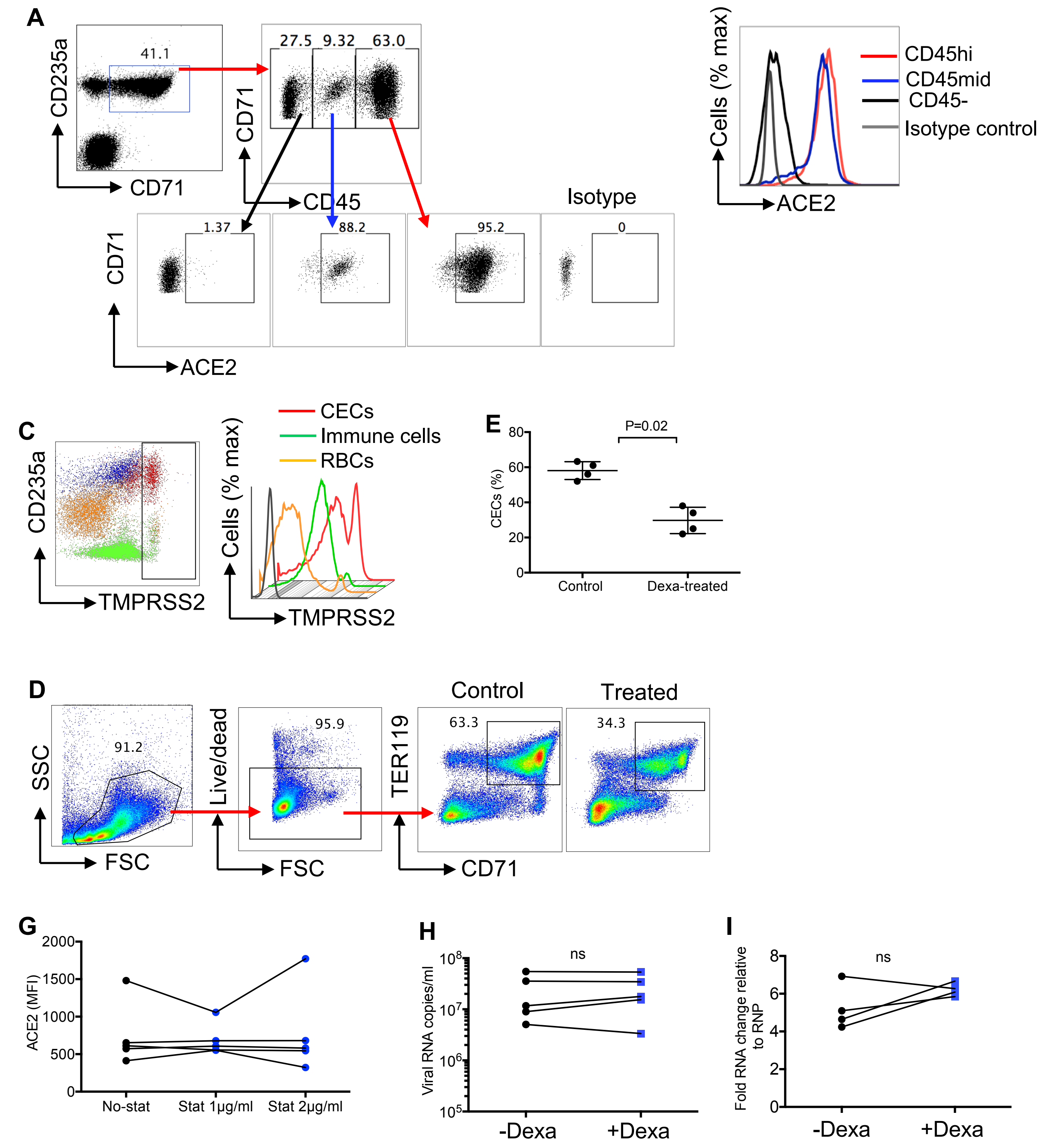

**Extended Data Fig Legends**

**Fig 1.** (**A**) Flow cytometry plots of gating strategy for CD71+CD235+ cells (CECs). (**B**) Representative flow cytometry plots, and (**C**) cumulative data of percentages of CD235a+CD71- cells in the PBMCs of COVID-19 patients versus healthy controls (HCs). (**D**) Representative plot of the expression of PDL-1 and VISTA on CECs of a COVID-19 patient. (**E**) Histogram of the purity of isolated CECs. (**F**) Cumulative data of CD107a in CD8 T cells following activation with SARS-CoV-2 peptides (2 μg/ml, 6 hr) without (W/out) or with CECs (**G**) Gating strategy for plasma cells. (**H**) Correlation of % antibody secreting cells with % CECs in PBMCs of COVID-19 patients. (**I**) The presence of ACE2 in different organs/tissues of mice are shown by western blot. (**I**) Representative flow plot of the purity of isolated CECs used for western blotting.

**Fig 2**. (**A**) Gating strategy for the identification of ACE2 expression on CECs, RBCs (CD235a+CD71-) and immune cell lineages (CD235a-CD71-). (**B**) Percentages of ACE2 expressing CECs among total CECs of a patient admitted to the ICU over time. (**C**) Representative, and (**D**) cumulative data showing intensity of CD147 on CECs versus RBCs (CD235a+CD71-) and other immune cell lineages as measured by mean fluorescence intensity (MFI). (**E**) Representative plots of CD26/CD147 co-expression on CECs versus RBCs (CD235a+CD71- cells). (**F**) Histogram of the intensity of CD26 expression on CECs and RBCs versus isotype control. (**G**) Representative flow plots showing expression of CD26 on CD45+CECs versus CD45-CECs.

**Fig 3**. (**A**) Quantification of soluble ACE2 by ELISA in the plasma samples of COVID-19 patients compared to healthy controls (HCs). (**B**) Comparing soluble ACE2 levels in the plasma of COVID-19 patients at the time of hospital admission and 5-7 days later. (**C**) Longitudinal quantification of soluble ACE2 in the plasma of 3 patients at day 3 until 43 days post the admission to the ICU. (**D**) Gating strategy for the identification of TMPRSS2 expression on CECs, RBCs (CD235a+CD71-) and immune cell lineages. (**E, F**) Representative gating strategy for the identification of TMPRSS2 expressing CECs using Image stream.

**Fig 4**. (**A**) The expression of TMPRSS2 in different organs/tissues of mice compared to the positive control (human colorectal adenocarcinoma grade II) are shown by western blot. (**B**) The expression of TMPRSS2 in CECs (CECs+) versus PBMCs-depleted of CECs (CECs-) from either COVID-19 patients or placental tissues (isolated CECs versus other cells (CECs-)) and cord blood compared to mouse lungs are shown by western blot. (**C**) Representative plots of TMPRSS2 surface expression on CD45+CECs of human bone marrow.

**Fig 5.** (**A**) Flow plots showing the % expression of ACE2 on CD45-, CD45lo and CD45hiCECs of the placental tissue compared to the isotype control. (**B**) Histogram of the intensity of ACE2 expression on CD45-, CD45lo and CD45hiCECs of the placental tissue compared to the isotype control. (**C**) Plots showing the intensity of the expression of TMPRSS2 on total CECs compared to immune cells lineages and RBCs of the placental tissue. (**D**) Gating strategy and representative plots of CECs (CD71+TER119+) in mice. (**E**) Cumulative data of % CECs in control versus treated mice with dexamethasone (1 μg/g body weight) and quantified 2 days later. (**F**) Plots showing expression of TMPRSS2 on CECs obtained from a COVID-19 patient once treated for 24 h with or without dexamethasone at 1 or 2 μg/ml. (**G**) Data of ACE2 expression on CECs obtained from COVID-19 patients once treated for 24 h with or without atorvastatin (stat) at 1 or 2 μg/ml as measured by mean fluorescence intensity (MFI). (**H**) Viral RNA copies in culture supernatants of monocytes either untreated or treated with dexamethasone (2 μg/ml) for 24 hr before infection with SARS-CoV-2 as measured 24 post infection. (**I**)

Cellular fold RNA changes relative to the housekeeping gene (RNP) in cell pellet of monocytes either untreated or treated with dexamethasone (2 μg/ml) for 24 hr before infection with SARS-CoV-2 as measured 24 hr post infection.
